# Spatial resource dynamics control resistance escape

**DOI:** 10.64898/2025.12.22.695823

**Authors:** Nico Appold, Timon Citak, Auguste Palm, Jona Kayser

## Abstract

The evolution of therapy resistance in structured populations such as biofilms and solid tumours is shaped by emergent spatial organization, with profound consequences for evolution-based therapies. However, how treatment reshapes these patterns remains poorly understood. Here we show that intermittent treatment pulses transiently reconfigure the resource landscape, reorganize spatial growth zones, and can enable resistant mutants to escape spatial confinement and drive therapy failure. We introduce a spatial evolution assay in which populations expand from single, genetically tailored yeast cells, enabling quantitative tracking of the full spatiotemporal trajectories of continually emerging resistant mutants under intermittent treatment. By integrating these observations with a mechanistically interpretable \textit{in silico} model in a real-to-sim-to-real loop, we identify an effective phase transition in schedule space that defines an optimal balance between population control and sustained resistance confinement, which we validate experimentally. Together, our results establish resource-mediated spatial confinement as a central organizing principle of resistance evolution and provide a mechanistic foundation for spatially informed, evolution-based therapies.

## 2 Main

Therapy resistance remains one of the biggest challenges in the treatment of pathological cellular populations, spanning bacterial infections, fungal pathogens, and cancer [1– Resistance-mediated therapy failure is driven by two tightly coupled processes: the continual *de novo* emergence of resistant mutants during population expansion and their subsequent amplification under treatment. In well-mixed, mean-field descriptions, these processes are often treated as separable, with mutations supplying resistant cells and therapy reshaping relative fitness. However, many clinically and ecologically relevant populations are neither well mixed nor exposed to spatially uniform environments. Biofilms, tissues, and solid tumours are typically compact, mechanically crowded, and structured by steep gradients of nutrients, oxygen, and signaling molecules. Such gradients can generate pronounced physiological heterogeneity, including growth-arrested subpopulations [6]. As a consequence, in structured populations resistance evolution becomes intrinsically spatio-temporal: Where a mutant arises within the population and when selective pressures change can be as important as the mutant phenotype.

Over the past two decades, a broad body of work has demonstrated that spatial population structure and emergent collective phenomena fundamentally reshape evolution in dense, range-expanding populations. For example, at expanding fronts, effective population sizes are small, competition is highly localized, and stochasticity is amplified [7–11]. Correlated founder effects can elevate mutants to large frequencies purely by positional advantage, while growth-induced collective motion can mask or renormalize fitness differences [12–14]. Spatial structure can inflate the frequency of mutational “jackpot” events [15], generate inflation-selection balances that stabilize rare subclones and drive evolutionary rescue [16], shape antagonistic interactions [17], and enhance conversional meltdown [18], among other effects. These studies increasingly capitalize on recent advances in molecular manipulation techniques specifically tailored for experimental evolution [19]. More recently, spatial transcriptomics and related spatial genomics approaches have revealed analogous patterns in solid tumours, underscoring that many of the same spatial-evolutionary principles inferred from microbial expansions also shape cancer evolution [20, 21]. Together, these studies establish evolution in compact, growing populations as an emergent process in proliferating active matter [22]. Because evolutionary activity is often concentrated near the leading edge of spatial population expansion, much of this work has focused on mutants originating at or near this front. More recently, attention has shifted toward the mutant-rich population bulk, which harbors many small, often undetectable lineages [23, 24]. While typically suppressed, these bulk-born mutants can become evolutionarily decisive when environmental conditions change.

Therapy onset represents such a drastic environmental change and thus provides a natural conceptual bridge between spatial evolutionary theory and the emerging paradigm of evolution-based cancer therapies. Rather than attempting to completely eradicate all cancer cells at the risk of accelerating resistance, evolution-based approaches deliberately exploit intrapopulation competition by maintaining a substantial sensitive population that suppresses resistant cells [25, 26]. So-called adaptive therapies, making closed-loop treatment adjustments based on population-level observables and mathematical evolution models, have shown promise in preclinical studies and early clinical trials [27–32]. Fixed-cycle intermittent schedules have been proposed as a minimal open-loop alternative that embodies a similar evolution-based rationale, but have so far shown limited success in clinical trials, potentially due to unmodeled heterogeneity in resistance phenotypes and schedule optimality [33–35]. In addition, spatially explicit simulations have demonstrated that local competition, spatial refugia, and heterogeneity in resource and drug fields can qualitatively alter tumour evolution and therapy outcomes compared to meanfield predictions [35–41].

Together, these advances highlight a growing convergence between spatial evolutionary theory and evolution-based therapy. Yet a quantitative, mechanistic understanding of how spatial heterogeneity, mutation supply, and time-varying selection jointly shape evolutionary dynamics and therapy outcome remains at large. In particular, we lack experimentally tractable systems that recapitulate the defining features of this scenario within a single, controllable platform: initiation from a single susceptible founder cell, range expansion into a densely packed population, continuous *de novo* mutation supply, and temporally structured therapy (Fig. 1a). Without such systems, it remains difficult to directly observe the full spatio-temporal trajectories of emerging resistant lineages, to rigorously test mechanistic hypotheses derived from theory and simulation, and to experimentally validate optimized treatment schedules in a spatially explicit setting.

**Figure 1:**
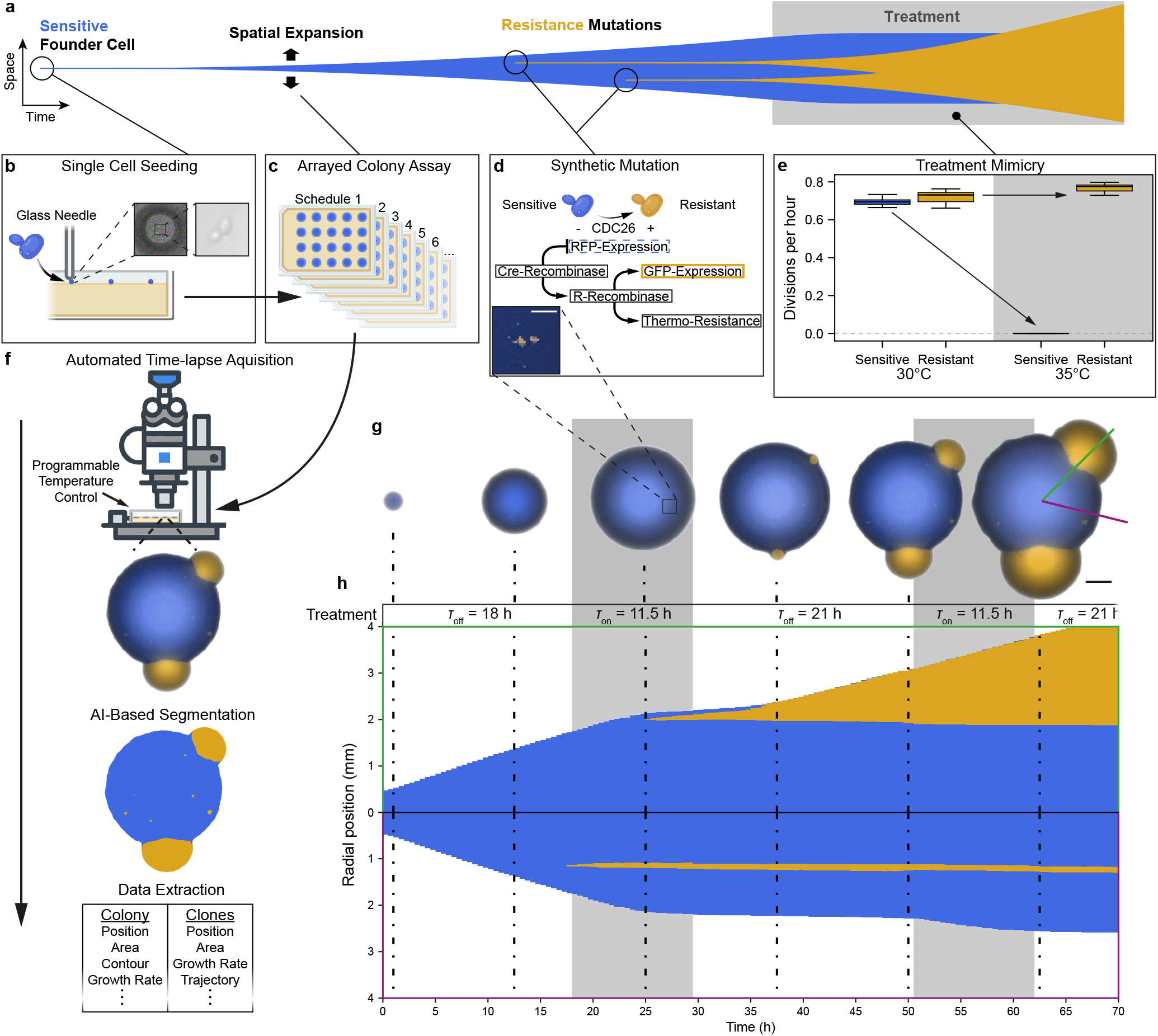
Experimental platform to study resistance under time-structured treatment. **a** Schematic of resistance evolution and therapy failure in spatially structured population. **b** Single cell seeding of sensitive cells. **c** Colony formation and time-lapse imaging under variable treatment schedules. **d** Synthetic mutation construct enabling stochastic, irreversible recombinase-mediated switching from thermo-sensitive to thermo-resistant cells coupled to a fluorescence color change (Supplementary Fig. 9). High magnification image of resistant clones within a colony. Scale bar, 100 µm. **e** Median growth rates of sensitive and resistant cells at 30°C and 35°C. Error bars indicate interquartile range (IQR), whiskers 1.5x IQR. n = 9 per experiment and cell type, with two replicates for 30 °C **f** Image analysis pipeline from automated temperature control and time-lapse imaging to segmentation and colony- and clone-level data extraction. **g** Representative intermittent-therapy time-lapse. Imaging starts at *t* = 0 h (24 h after seeding). Resistant cells (yellow) expand while sensitive cells (blue) are arrested. Time points: 1, 12.5, 25, 37.5, 50 and 62.5 h after imaging start. Scale bar, 1 mm. **h** Radial kymographs from the colony in **g** (colored radials), showing escape (top, 104°) and confinement (bottom, 46°) of bulk-born resistant mutants. Gray shading indicates treatment. Dash-dotted lines mark time points in panel e.

Here, we introduce a high-throughput microbial therapy-mimicry assay that enables direct tracking of the full spatio-temporal trajectories of continually emerging resistant clones in range-expanding yeast populations under programmable treatment schedules. Coupling these experiments to a mechanistically interpretable spatial digital twin reveals otherwise hidden resource landscape dynamics. Leveraging this integrated experimental and mechanistic modeling framework, we uncover therapy-induced reconfiguration of the spatial resource landscape as the key mechanism governing confinement, release, and reconfinement of resistant mutants. This mechanism gives rise to a sharp boundary in intermittent therapy space that separates sensitive-dominated control from resistance-driven failure and simultaneously defines an optimal therapeutic balance. Closing the real-to-sim-to-real loop, we experimentally validate this predicted sweet spot.

## 3 Results

### 3.1 A platform for tracking resistant mutants under time-modulated therapy

To quantitatively probe how *de novo* resistance emergence interacts with time-modulated, therapy-driven population dynamics in a spatially structured setting, we designed an experimental evolution assay based initiated from individually placed, genetically tailored *S. cerevisiae* cells (Fig. 1b). These founder cells expand into compact populations in a high-throughput arrayed colony assay (Fig. 1c).

To generate a continual supply of trackable resistant mutants, we engineered a stochastic and irreversible *synthetic mutation* system that couples resistance to a change in fluorescence color via a hierarchical dual-recombinase switch (Fig. 1d, Supplementary Fig. 9a). Time-dependent treatment is implemented via programmable temperature shifts, allowing precise control over therapy onset, duration, and pause (Fig. 1e, Supplementary Fig. 9b). In the unmutated state, cells are thermo-sensitive and proliferate at 30°C but arrest at 35°C, whereas mutated cells become thermo-resistant and continue to proliferate at elevated temperature. Because this mutation is irreversible and fluorescently reported, resistant lineages can be identified and tracked throughout colony growth.

We combine this assay with multi-day time-lapse fluorescence microscopy and automated temperature control to record population histories for up to 160 h (> 100 generations) (Fig. 1f). To quantitatively extract colony-and clone-level dynamics across the entire population history, we developed a custom image analysis pipeline combining automated acquisition with machine-learning–based segmentation.

Fig. 1g shows a representative time series under an intermittent schedule. To visualize spatio-temporal dynamics along the direction of expansion, we compute radial kymographs from selected colony radials (Fig. 1h; colored radials in Fig. 1g). We observe two qualitatively distinct fates for resistant clones. While some can permanently escape confinement as continually growing domes, most resistant clones remain confined within the population bulk.

Together, these observations establish a baseline phenomenology: in spatially structured, range-expanding populations, de novo resistant mutants are continually generated but are typically confined within the bulk. Therapy can alter this fate, enabling some bulk-born resistant clones to escape, while others remain suppressed. In the following sections, we investigate the mechanisms governing this confinement and release, and how they can be controlled by the timing of therapy.

### 3.2 Pausing therapy spatially reconfines resistant mutants

To systematically study therapy response dynamics, we first examined mutant emergence and expansion under continuous therapy (CT) (Fig. 2a). Colonies were expanded at 30 °C (therapy off) and then shifted to 35 °C (therapy on), where temperature was held constant for the remainder of the experiment.

**Figure 2:**
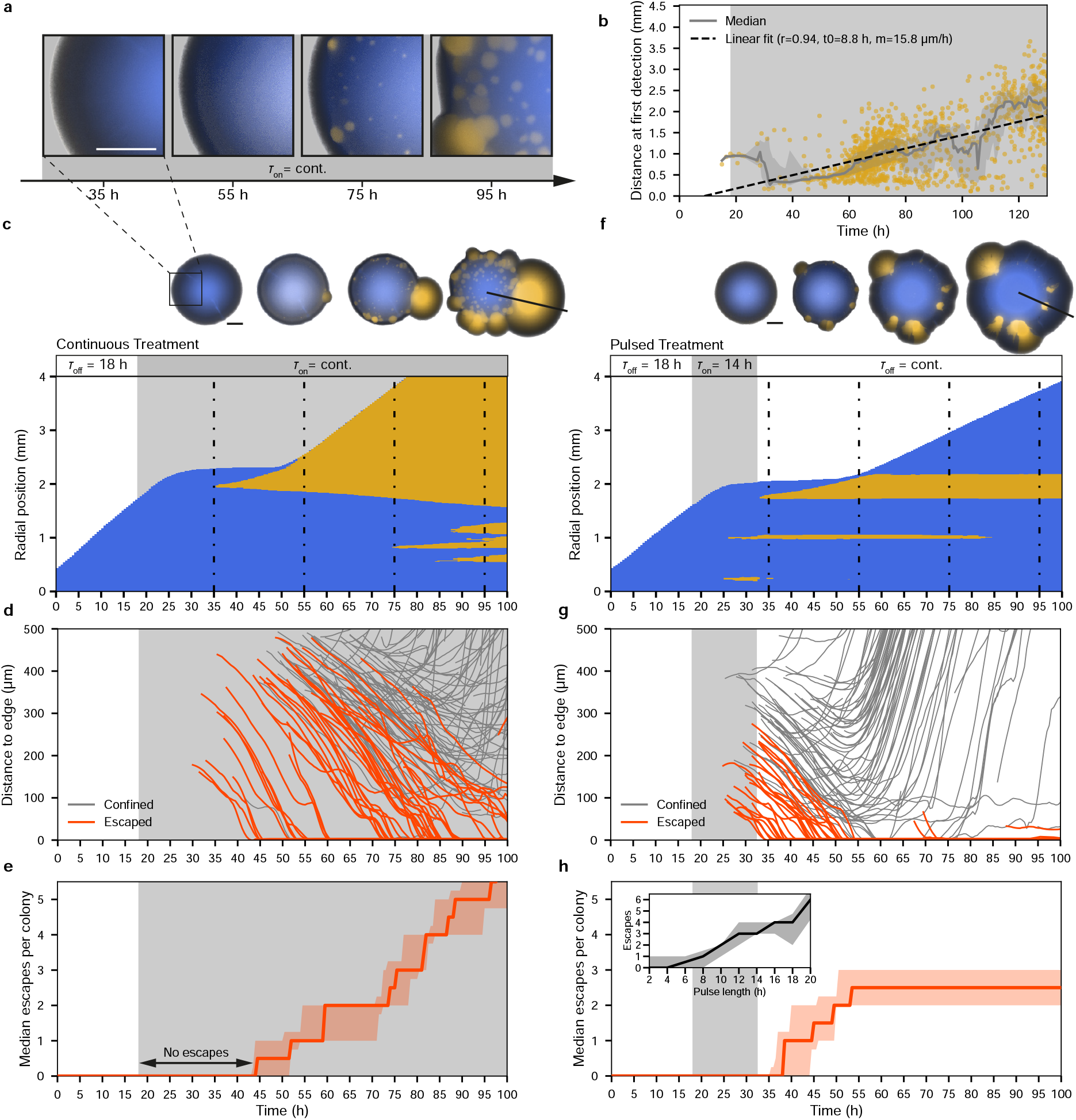
Confinement of resistant clones under pulsed therapy. **a** High magnification time-lapse images showing expansion of resistant clones during treatment. **b** Distance of resistant clones to the colony front at first detection during continuous therapy (median ± interquartile range). Dotted line shows a ReLu fit to all data points. **c, f** Radial kymographs with corresponding time-lapse images for representative colonies under continuous (c) and pulsed (f) schedules. Dash-dotted lines mark shown time points. Scale bars, 1 mm **d, g** Trajectories of the front position of individual resistant clones over time for all colonies under continuous (panel d, *n*_colonies_ = 9) and pulsed (panel g, *n*_colonies_ = 12) treatment. Clones were included if first detected after 25 hours, prior to proliferation arrest of sensitive cells. Clones were classified as escaped if their distance to the colony front did not increase after approaching within 20 µm. **e, h** Median cumulative number of escaped clones per colony under continuous and pulsed treatment (panel e, h). Inset, escape events per colony across pulse durations (median ± IQR). **a-h** Gray shading indicates treatment.

Despite the spatial homogeneity of the temperature shift, mutant expansion exhibits a pronounced spatio-temporal structure: the time at which a resistant mutant is first detected is strongly correlated with its distance from the colony front (Fig. 2b). Thus, resistant lineages do not become detectable throughout the colony simultaneously upon therapy onset, but rather appear progressively as a function of position, indicating that mutant growth is gated by a spatially varying growth potential within the colony.

Tracking individual mutant trajectories under CT reveals a characteristic approach toward the colony edge until permanent escape occurs (Fig. 2c, d). Notably, mutants originating further inward can sometimes remain confined when a more forward mutant lineage occupies the advancing front. At the colony level, these dynamics imply a finite time window after therapy onset during which no mutant has yet escaped; as therapy continues, this window closes and the cumulative number of escaped mutants increases steadily (Fig. 2e).

The existence of a delay between therapy onset and resistance escape suggests that a sufficiently short therapy pulse could be used to transiently suppress overall population expansion while still preventing mutants from reaching the colony edge. Moreover, the observation of local clonal interference raises the possibility that restoring sensitive population growth could act as an effective barrier at the front, enforcing reconfinement of resistant mutants.

To test this idea, we applied therapy as before but interrupted treatment after a pulse duration of *τ*_on_ = 14 h, allowing sensitive cells to resume growth (Fig. 2f). At the time of treatment interruption, no resistant mutant had yet escaped. Strikingly, this therapy pause indeed restored spatial confinement for many clones: mutants that had expanded during the pulse ultimately stalled and remained confined within the colony bulk without escaping (gray traces in Fig. 2g). Notably, the time at which these lineages became re-confined, as quantified by their closest approach to the colony front, often preceded the full recovery of sensitive population expansion, consistent with the re-establishment of an effective competitive barrier by sensitive cells. Nonetheless, because sensitive growth resumed only after a substantial regrowth lag, mutants residing sufficiently close to the front could continue advancing and escape for a limited period after therapy cessation (Fig. 2f-h).

Systematically varying the pulse duration shows that shortening treatment pulses strongly enhances spatial confinement (Fig. 2h, inset). In particular, for pulses shorter than *τ*_on_ < 6 h, resistance escape is almost completely suppressed, indicating that the maximum pulse length that reliably prevents escape is more restrictive than what would be inferred from CT trajectories alone, due to the additional time available for mutant advance during the post-pulse regrowth lag.

A possible explanation for these observations is that therapy reshapes the spatial resource landscape that naturally arises in many spatial population expansions [9, 13, 15]. More specifically, suppressing sensitive growth permits nutrients to redistribute from the colony periphery toward the interior, enabling bulk-born mutants to expand, whereas renewed sensitive growth restores resource depletion in the bulk and counteracts mutant advance. Nutrients are therefore a likely candidate for the hidden spatial field that gates mutant expansion, but directly measuring their spatio-temporal dynamics under time-modulated therapy is experimentally challenging. We therefore next introduce a mechanistically minimal digital twin that couples growth to a diffusing nutrient field in a spatial setting to test whether the observed release and reconfinement dynamics emerge naturally from this coupling.

### 3.3 An *in silico* model couples spatial resource dynamics to resistance escape

To test whether therapy-induced release and reconfinement can emerge generically from coupling population growth to a spatial nutrient field, we constructed a stochastic simulation framework as a mechanistically minimal “digital twin” in which nutrient dynamics are directly accessible (Fig. 3a). The model integrates nutrient-dependent proliferation, growth-coupled spreading, and nutrient diffusion in two spatial dimensions. Growth and dispersal are spatially coupled: local proliferation drives outward spreading at the colony periphery, and this front motion continuously relocates the growth zone toward regions of higher nutrient availability similar to a travelling wave. As a result, population expansion is not merely enabled by nutrients, but also reshapes nutrient access by shifting where consumption occurs in space.

**Figure 3:**
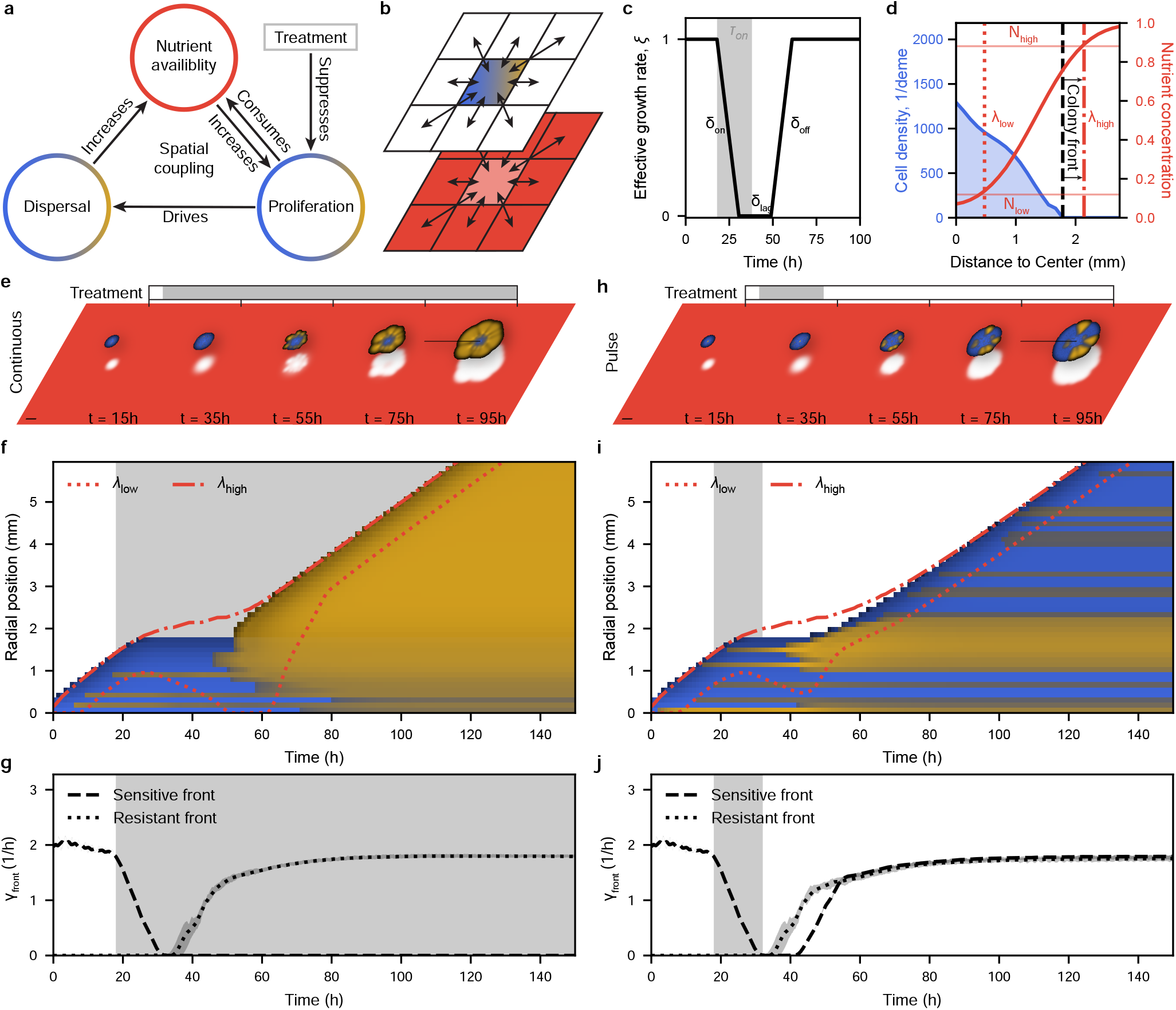
Simulation of spatial resource dynamics. **a** Schematic of spatial couplings between nutrients, proliferation, dispersal, and treatment. **b** The simulation consists of sensitive (blue) and resistant (yellow) populations interacting with each other via an underlying nutrient landscape (red), organized on a 2D square lattice. **c** Treatment modifies growth rates with delays, leading to an effective growth rate *ξ*. The specific form of the modification is defined by 3 parameters: *δ*_on_, the time it takes for treatment to take full effect, *δ*_lag_, the time *ξ* stays constant after treatment ends, and *δ*_off_, the time it takes until *ξ* fully recovers. In case of treatment stopping before *ξ* reaches zero an additional overshoot phase *δ*_over_ occurs, which is part of *δ*_lag_. Gray shading indicates treatment. **d** The nutrient dynamics can be visualized via a lower nutrient boundary λ_low_, defined by a set nutrient threshold and an upper nutrient boundary λ_high_ (methods). **e, h** Time lapse of a continuous therapy (CT) and a 14 h pulse simulation run. The black line marks the position of the kymograph in panel f, i. Scale bar indicates 2 mm. Opacity indicates density. Sensitive and resistant cells are displayed with gamma correction *γ* = 5 and *γ* = 2.5, respectively. **f, i** Kymograph for the continuous dose and pulse treatments in panel e, h. The red lines show the nutrient dynamics as defined in panel d. Gray shading indicates treatment. The colors are displayed with gamma correction *γ* = 10 for all types. **g, j** Median average front cell growth speed *γ*_front_ of sensitive and resistant cells (n = 20). The shaded area indicates the IQR. Gray shading indicates treatment.

In short, space is discretized into a 2D grid of demes that contain sensitive cells *S* and resistant cells *R*. Cells proliferate at a nutrient-dependent rate and undergo growth-driven local spreading (Fig. 3b). Resistant mutants arise stochastically during sensitive growth, such that mutation supply is coupled to proliferation as in our experiments. Therapy is implemented as a modulation of sensitive growth by a factor *ξ* ∈ [0, 1] that captures experimentally observed response kinetics, including an on-delay, a post-treatment lag, and an off-delay (Fig. 3c). Importantly, competition is mediated solely through nutrient limitation (no explicit local carrying capacity), so that both cell types interact only indirectly via a shared nutrient field. Nutrients are modeled as a freely diffusing consumable resource that is depleted by local growth (methods).

To visualize the shape of the nutrient gradient and its therapy-induced changes, we compute the position of two complementary nutrient thresholds: the low-threshold position λ_low_ and the high-threshold position λ_high_ (Fig. 3d). Whereas λ_low_ reports how far growth-permitting nutrients penetrate into the bulk, λ_high_ reports how tightly the leading edge remains coupled to the high-nutrient side of the gradient.

Using the parameterized model, we simulated population trajectories under continuous therapy and under a finite pulse of *τ*_on_ = 14 h (Fig. 3e,h). Despite its minimal structure, the digital twin qualitatively reproduces the key spatio-temporal phenotypes observed experimentally: resistant mutants originating in the bulk can become released and advance toward the colony edge under therapy, while pausing therapy can prevent escape and restore spatial confinement in a duration-dependent manner.

Because nutrients are explicitly modeled, we can directly link these behaviors to therapy-induced dynamics of the growth-permitting layer. Under continuous therapy, suppressed sensitive growth reduces nutrient consumption, allowing nutrients to penetrate inward and causing λ_low_ to move inward over time (Fig. 3f). This progressive expansion of the growth-permitting region releases bulk-born resistant mutants and ultimately enables them to reach and escape at the colony edge. Fig. 3g shows the type-specific growth rate *γ*_front_ of cells at the front.

Under a finite pulse, by contrast, both the increase of λ_low_ and reduction in *γ*_front_ are transient: if therapy is halted sufficiently early for *γ*_front_ to recover, escape is reduced and the nutrient landscape relaxes toward its pre-treatment state, restoring spatial confinement (Fig. 3h-j).

Together, Fig. 3 supports a simple mechanistic picture consistent with the experimental observations: spatial growth and nutrient consumption generate a nutrient gradient along the expansion direction, and growth-driven front advance continuously couples the colony edge to the high-nutrient side of this gradient. Therapy transiently weakens this coupling by suppressing sensitive growth, broadening the peripheral nutrient landscape and releasing bulk-born resistant mutants. Interrupting therapy can prevent escape by reforming a competitive barrier of sensitive cells at the colony edge and restore confinement by reestablishing nutrient depletion in the bulk.

### 3.4 An effective phase transition in intermittent schedule space defines a therapeutic sweet spot

Having identified the mechanisms governing mutant release and reconfinement, we next return to therapy optimization which poses an intrinsic tradeoff: suppressing sensitive growth slows overall colony expansion but increases the risk of resistant mutant escape; conversely, allowing sensitive growth helps to maintain spatial confinement, but comes at the cost of continued population expansion. Aiming to balance growth suppression with confinement naturally motivates a concatenation of therapy-on and therapy-off intervals, often referred to as intermittent therapy (IT). However, the observed spatial couplings render a straight-forward identification of optimal combinations of interval lengths *τ*_on_ and *τ*_off_ elusive.

We used the digital twin to scan a large space of schedules ***τ*** = (*τ*_on_, *τ*_off_) (Fig. 4a,b; Supplementary Fig. 6). We quantify schedule performance by the time to progression (TTP), defined as the time required for colony area to progress above a predefined threshold (methods). Across schedule space, TTP exhibits a pronounced ridge of optimal schedules, ***τ*** ^∗^, separating two modes of early progression (Fig. 4a). On one side, progression is sensitive-dominated (undertreatment); on the other, resistant-dominated (overtreatment), with a sharp switch in resistant fraction near ***τ*** ^∗^ (Fig. 4b). Thus, ***τ*** ^∗^ marks a narrow line of “sweet spots” that maximizes delay while maintaining confinement.

**Figure 4:**
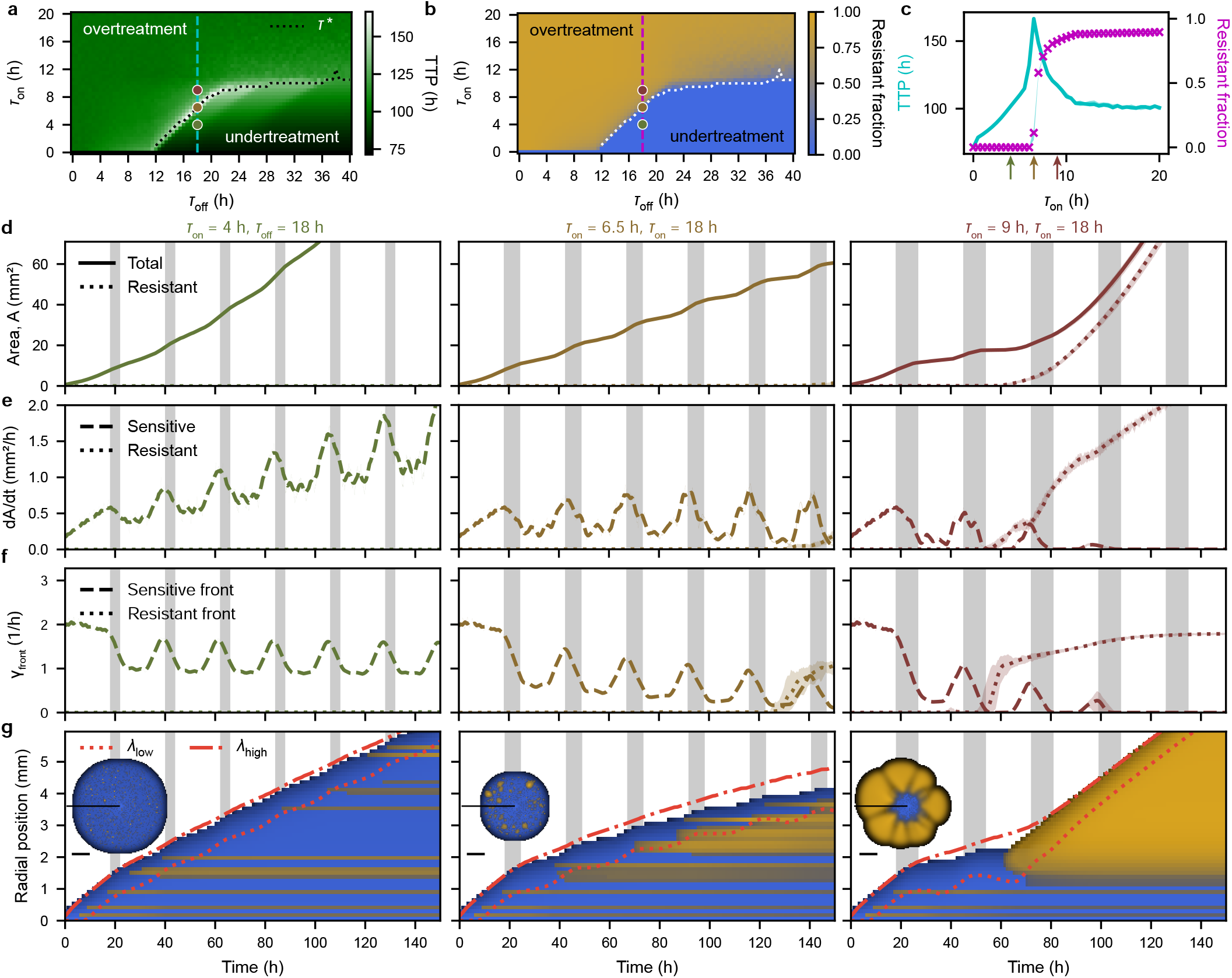
An effective phase transition in treatment space defines therapy outcome. **a** Parameter sweep over treatment on *τ*_on_ and off *τ*_off_ pairs showing the median time to progression (TTP, see main text) (n = 20 runs for each *τ*_on_, *τ*_off_ pair). Colored dots mark the schedules shown in panel d-f. The dashed vertical line correspond to the TTP profile shown in panel c. The dotted line shows the maximum TTP position for each *τ*_off_ value and is defined as *τ* ^∗^. **b** Median resistant fraction (n = 20) for the sweep shown in panel a. The dashed vertical line correspond to the resistant fraction profile shown in panel c. **c** Median TTP and resistant fraction profiles together with their respective interquartile ranges (IQR) for the dashed vertical lines in panel a, b. IQR are to small to be displayed (Supplementary Fig. 6). **d - f** Median area A, median area growth rates d*A*/d*t*, and median average front cell growth speed *γ*_front_ trajectories for the treatment schedules *τ*_on_ = 4 h, 6.5 h, 9 h and *τ*_off_ = 18 h. The shaded area is the interquartile range (n = 20). Gray shading indicates treatment. **g** Kymograph together with the nutrient dynamics indicated by the red lines for one example angle and simulation run of the above schedules. The colony image shows the entire example colony at 120 h with the scale bar indicating 2 mm. The black line shows the position of the kymograph in the colony. Gray shading indicates treatment. In the colony images sensitive and resistant cells are displayed with gamma correction *γ* = 5 and *γ* = 2.5, respectively, while in the kymograph colors are displayed with gamma correction *γ* = 10 for all types.

To connect this macroscopic optimum to within-cycle dynamics, we examined a representative slice at *τ*_off_ = 18 h (Fig. 4c) and three schedules spanning the optimum (Fig. 4d-g; *τ*_on_ = 4 h, 6.5 h, 9 h). For short pulses, *γ*_front_ rebounds during each pause, rapidly restoring a sensitive rim at the edge. This reinstates bulk depletion and allows λ_low_ to retreat close to its pre-pulse value, rendering consecutive cycles approximately independent. For 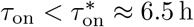, this leads to an almost constant cycle-averaged growth. Beyond 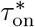, by contrast, therapy depresses *γ*_front_ strongly enough that front recovery during *τ*_off_ is incomplete. The resulting cycle-to-cycle drift of λ_low_ produces a ratchet-like loss of confinement, explaining the sharp rise in resistant fraction and the sensitivity of outcomes to minute schedule changes (Fig. 4f, g). Consistent with this picture, the linear increase of ***τ*** ^∗^ for intermediate pauses follows from therapy response kinetics, whereas for long pauses the boundary is set by single-pulse escape and for very short pauses by the post-treatment regrowth lag.

The sharp switch between a history-independent (approximately Markovian) regime and a history-dependent (non-Markovian) regime is reminiscent of a phase transition in schedule space. Although our system is far from equilibrium, this analogy provides a useful conceptual framework: optimal schedules lie near the boundary where front recovery remains barely sufficient to maintain reconfinement, making the “sweet spot” narrow and sensitive to parameter changes.

Finally, we do not expect the simplified digital twin to quantitatively reproduce all experimental details. In particular, the precise location of ***τ*** ^∗^ is sensitive to the shape and magnitude of therapy-response delays that govern *γ*_front_ (Supplementary Fig. 5). Quantitative predictions from the sweep should therefore be interpreted as rough guides. Nevertheless, the underlying mechanism is generic and should remain relevant in more complex experimental settings.

### 3.5 Experimental validation of optimal intermittent therapy

Guided by the digital twin, we tested the observed principles empirically by applying the same set of schedules (*τ*_off_ = 18 h; *τ*_on_ = 4 h, 6.5 h, 9 h) in experiments, in addition to no-treatment and continuous treatment references (Fig. 5a).

Comparing overall therapy performance by measuring the time to progression (TTP) and resistant fraction, we find a characteristic optimality very similar to that observed in the digital twin (Fig. 5b, c). TTP initially increases with longer treatment duration but then peaks at *τ*_on_ = 6.5 h (Fig. 5b). This peak also correlates with a sudden rise in mutant abundance (Fig. 5c).

**Figure 5:**
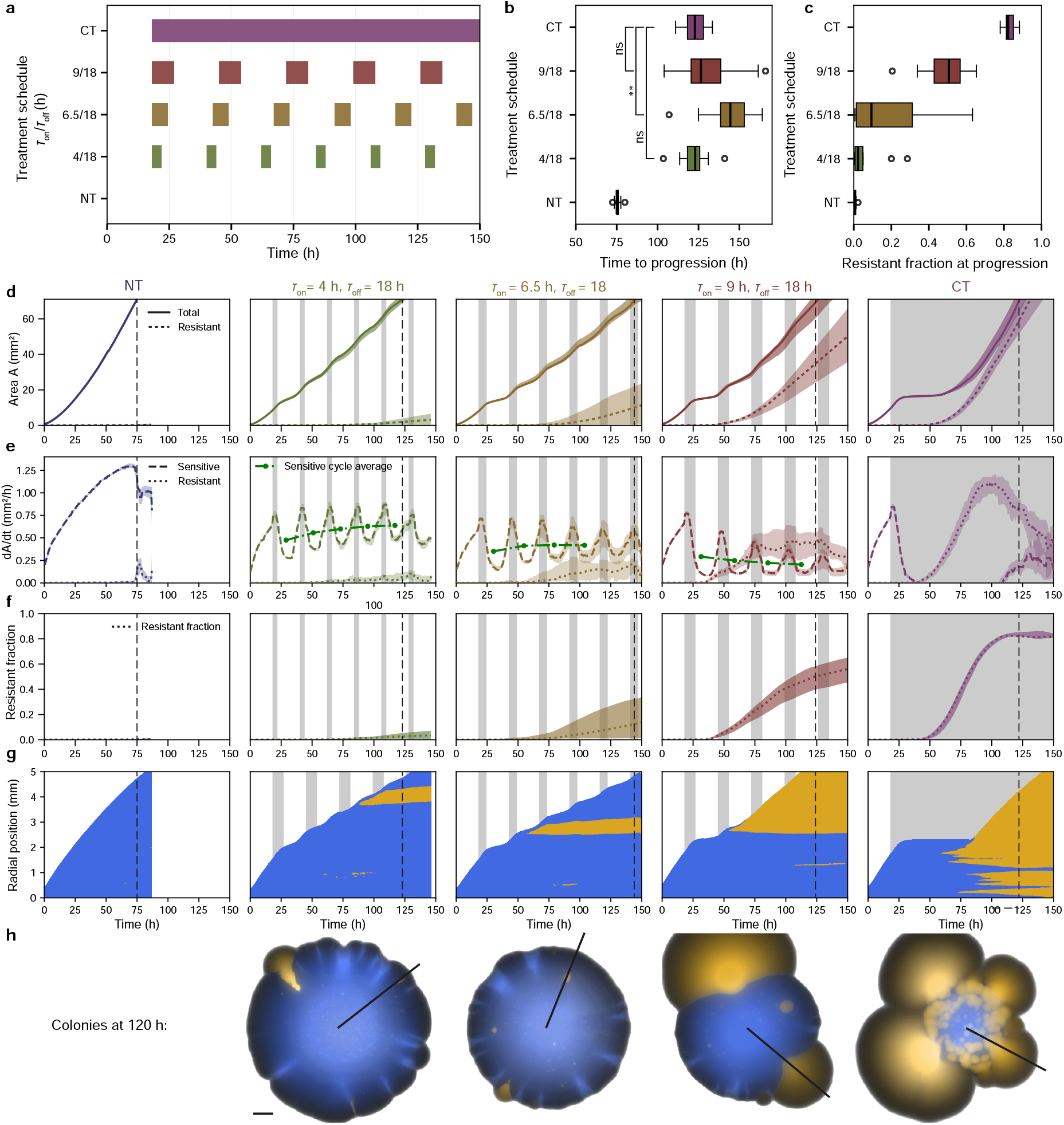
Experiments support delayed progression found in simulated treatment schedules. **a** Experimentally tested schedules and corresponding *τ*_on_ intervals. **b** Time to progression (TTP), defined as the first time point at which colony area reached ≥ 71 mm^2^, for indicated treatment schedules. Boxplots show median, IQR; whiskers, 1.5x IQR. *P* values correspond to one-sided Mann-Whitney U-tests with correction (*P* = 0.0079 for 6.5 h / 18 h). n values are indicated in table. 1 **c**Resistant cell fraction at TTP (median, IQR; whiskers, 1.5x IQR) **d** Median total and resistant colony areas for no treatment (NT), intermittent schedules (*τ*_on_*/τ*_off_ : 4 h / 18 h, 6.5 h / 18 h, 9 h / 18 h), and continuous treatment (CT). **e** Median growth dynamics of sensitive and resistant populations, including cycle-averaged sensitive growth. **f** Median temporal evolution of the resistant fraction (median, IQR). **g** Radial kymographs of representative colonies for *τ*_on_/*τ*_off_ : 4 h / 18 h, 6.5 h / 18 h, 9 h/ 18 h and CT (angles 52°, 22°, 130° and 118°. **h** Images at 120 h corresponding to g; black radial marks the sampled angle. Scale bar, 1 mm. In panel d-f dashed black lines indicate median TTP. Grey shading denotes treatment periods

Measuring how the area of sensitive and resistant cell population changes over time, analogous to the analysis of simulation data in Fig. 4, we find a behavior that qualitatively matches that observed in the digital twin (Fig. 5d-g). For short treatment durations (*τ*_on_ = 4 h), therapy induced growth rate reductions of sensitive cells are balanced by a rebound of growth during therapy pauses, resulting in a slowly increasing cycle-averaged growth rate and strong confinement of resistant mutants (Fig. 5e, f). For *τ*_on_ = 6.5 h, the cycle-averaged growth rate remains almost constant throughout the experiment, paired with a noticeable but still small increase in resistance fraction. Intriguingly, with *τ*_on_ = 9 h, experiments indeed exhibit the cumulative decrease in sensitive cell growth and the surge in mutant escapes that is characteristic for the resistance dominated region of schedule space (Fig. 5e, f; *τ*_on_ = 9 h).

While these empirical results only cover a very small, albeit critical, part of the therapy space, they are strong evidence that the concept of a therapy dependent phase transition also applies to the much more complex experimental realm. Moreover, they are a direct demonstration that the intrinsic coupling between spatial population structure and a therapy-modulated nutrient landscape can be exploited for therapy optimization while concomitantly making it challenging to hit the therapeutic sweet spot of optimal balance. However, the pre-defined regular intervals used here to explore the fundamental mech-anisms are just a thin slice of the large space of all possible schedules. For example, we could further improve therapy by making closed-loop adaptive treatment decisions based on mutant position, an avenue that warrants further investigation (Supplementary Fig. 10).

## 4 Discussion

This work shows that in structured populations, resource fluxes are a prime determinant of spatial growth patterns. Therapy can transiently disrupt these patterns, resulting in the escape of previously confined resistant mutants from the population bulk.

To systematically study the processes that govern these dynamics, this work introduces a therapy mimicry assay based on a genetically tailored yeast model. Pairing empirical results with an experiment-parameterized digital twin in a real-to-sim-to-real approach, we probe a large therapy design space, identify an intermittent therapy schedule that optimally balances population growth and resistance confinement, and then verify this *in silico*-derived strategy in experiments.

Three main conceptual insights emerge from this study: First, in populations undergoing resourcelimited range expansion, therapy-induced halt of peripheral growth intrinsically results in a redistribution of resources towards the deprived bulk, which can release previously confined resistant mutants from resource-mediated confinement. Second, strategically pausing treatment can be leveraged as a general control principle by restablishing resource-mediated confinement via rebounding sensitive cell growth at the at the population front. Third, small changes in treatment duration can cause dramatic changes in therapy outcome by switching from a Markovian to a non-Markovian treatment response class. The resulting effective phase transition defines a narrow therapeutic sweet spot that is characterized by treatment response kinetics.

The fundamental nature of the uncovered mechanisms suggests that similar effects may be present in a much wider class of spatially structured populations with resource gradients, including many bacterial biofilms and solid tumours. A broad collection of previous work has shown how emergent spatial and physical phenomena reshape the evolutionary dynamics at the expanding front of these systems[9, 11, 13–16, 37, 38]. Our work builds on these insights by showing that shifting fitness landscapes, with modulated therapy being an extreme example, can transiently shift growth towards the mutant-rich population bulk, drastically increasing the genetic diversity accessible to evolution. Importantly, strategically controlling intrapopulation competition is foundational to the new paradigm of evolution-based tumour therapies[25, 28, 39, 40]. In this context, our findings offer a new perspective on how the inherent resource gradients in solid tumours may be optimally exploited for improved therapy design. For example, using growth rebound as a clinically accessible measure for confinement potential in addition to overall burden might offer valuable guidance for treatment decisions. In addition, poorly vascularized tumour types, often associated with steep gradients and resource-depleted core regions, appear as ideal candidates for evolution-based therapies. Complementary treatments targeting angiogenesis, such as VEGF inhibitors, may further enhance the effectiveness of such therapy strategies. In addition, our results may offer an additional rationale explaining why recent clinical trials of intermittent therapy did not improve progression-free survival in patients, despite strong preclinical evidence[34, 35]. Finding the narrow therapeutic sweet spot is inherently challenging due to the biphasic nature of the therapy space while the commonly applied lead-in phase may eradicate it altogether.

While the effects observed in our well-controlled microbial model system might be general in nature, there will be additional influences on the evolution of resistant subclones in more complex systems. For example, both bacteria and tumour cancer cells actively migrate and interact with each other in a complex 3D microenvironment, which is absent in our models. In addition, our work primarily discusses therapy trajectories for individual populations, while systemic therapies, including evolution-based tumour therapy, is most relevant in a metastatic scenario with multiple heterogeneous lesion sites. It should also be noted that for both, antibiotic and anti cancer therapies, treatment typically causes cell death rather than the reversible inhibition of growth studied here. It will be therefore quintessential to additionally test the nuances of resource-mediated spatial confinement and derived therapy strategies in more realistic model systems that better capture these complexities, such as cancer cell organoids or biofilms of matrix-producing bacteria. Importantly, while the highly reproducible growth dynamics of our well-controlled model systems are ideally suited to study fundamental mechanisms via predefined intermittent schedules, real tumours exhibit a wide bandwidth of growth parameters, requiring more adaptive approaches.

In conclusion, our work underscores the critical role of an inherent spatio-temporal coupling between therapy-modulated nutrient landscapes, mutant growth and their escape in densely packed cellular populations. In essence, therapy failure risk is governed by whether therapy shifts the resource gradient long enough for bulk-born mutants to permanently escape confinement. Integrating the gained insights into our understanding of resistant evolution and therapy response will open new avenues for combating resistance-mediated therapy failure.

## 5 Methods

### 5.1 Strains

All experiments were performed using the genetically engineered strain yNA16 of the budding yeast *Saccharomyces cerevisiae*. yNA16 is derived from the ancestor strain yJK19 and carries a β-estradiol-inducible Cre-EBD recombinase expressed from the *SCW11* promoter and integrated at the *HO* locus. yNA16 harbors a synthetic, irreversible recombination-based mutation cassette (*PGPD-loxP-yEmRFP-ubq-cyh2r-TCYC1-kanMX-loxP-yEGFP-ubq-Rrec*), enabling stochastic switching from a sensitive to a resistant state accompanied by a fluorescent color change.

Thermosensitivity was introduced via an auxotrophic insertion at the *CDC26* locus (*pCDC26-RS-URA3-tURA3-RS-CDC26*). To enhance fluorescent signal in the switched state, a second activatable GFP cassette (*pGPD-RS-LEU2-tLEU2-RS-yEGFP-tADH*) was integrated. A switched derivative strain (yNA16s) was generated by inducing Cre activity with 3 µM β-estradiol in liquid culture, followed by selection at 35 °C. A resistant clone exhibiting stable green fluorescence was isolated and used for down-stream experiments.

### 5.2 Construction of yNA16

yNA16 was generated through sequential genetic modifications of yJK19 [16]. First, the Cre-responsive switching cassette was integrated to generate yNA14. Thermosensitivity was subsequently introduced at the *CDC26* locus, yielding yNA15. Finally, an additional recombination-activated GFP cassette was inserted to improve fluorescence intensity in the switched state, resulting in yNA16. All genetic constructs were amplified by PCR and assembled using NEBuilder HiFi DNA Assembly (New England Biolabs). The underlying concepts were inspired by previous works [14, 15, 18, 42].

### 5.3 Yeast Transformation

Yeast transformation were performed using lithium acetate/polyethylene glycol (LiAc/PEG)-based protocol. Yeast strains were inoculated into 5 ml YPD and grown overnight at 30 °C. The follwoing day, cultures were diluted into 50 ml YPD to an initial *OD*_600_ of 0.2 and grown at 30 °C with shaking for 3-4 h until reaching *OD*_600_ of ≈ 1.0. Cells were harvested by centrifugation (300 x g, 5 min), washed once with sterile distilled water, and resuspended in 1 ml distilled water. Aliquots of 100 µl were pelleted and resuspended in a transformation mix containing 50 % (w/v) PEG 3500, 1.0 M lithium acetate, boiled single-stranded carrier DNA, and either PCR product DNA or plasmid DNA. Cells were incubated at 42 °C for 40 min, pelleted, and gently resuspended in 250 µl YPD. Transformants were plated on YPD agar or selective media and incubated at 30 °C. For antibiotic selection, colonies were replica-palted onto YPD plates containing the corresponding antibiotic after overnight recovery.

### 5.4 Measuring growth rate

Growth rates of yNA16 and yNA16s at 30 °C and 35 °C were determined using plate reader assays. Thin liquid cultures were diluted 1:1000, and 10 µl of each culture was added to 450 µl of YPD per well in a 48-well plate. Each strain was measured in 16 replicates per plate, with *OD*_600_ recorded every 10 min. Plates were continuously shaken during incubation to promote exponential growth of all cells in suspension. Doubling times were estimated by fitting an exponential curve to the growth trajectories. The experiment at 30 °C was repeated twice to account for stochastic variation.

### 5.5 Arrayed colony assay

For the standard arrayed colony assay, *yNA16* populations were pre-cultured on YPD + 2 % agar supplemented with 200 µg/ml G418 for 2 days, then stored at 4 °C for up to 3 weeks. Monoclonal colonies from these plates were used to inoculate 3 ml liquid YPD medium containing 200 µg/ml G418 and grown for approximately 5 h at 30 °C. Cultures were vortexed for 5 min to disperse cell clusters, and 30 µl were spread in a narrow band on one side of a monowell plate containing 70 ml of 1 % agar YPD. The Plates were dried under sterile conditions with the lid open for 45 min prior.

Single cells, preferentially with a small bud, were isolated using a Singer MSM400 dissection microscope and placed in a 5 × 4 grid, yielding a potential maximum of 20 colonies per plate. Plates were incubated at 30 °C and 85 % relative humidity for 24 h to allow initial colony expansion, after which they were transferred to an on-stage incubation chamber for time-lapse imaging. Multichannel images were acquired every 30 min for up to 7 days. Temperature-based treatments and therapy holidays were implemented by adjusting the incubation chamber temperature manually or via a pre-programmed script. A type K thermocouple, placed on the agar surface at colony height, was used to monitor on-plate temperature. Colonies displaying growth defects, petite mutations, contamination, or physical obstruction due to agar irregularities were excluded from analysis. A summary of all experiments and the corresponding numbers of analysed colonies is provided in Table 1

**Table 1:** Experimental setups and corresponding number of colonies.

| Name or Schedule ($\tau_{\text{on}}/\tau_{\text{off}}$) | No. Colonies |
| --- | --- |
| No treatment | 13 |
| Continuous treatment | 9 |
| Pulse (14 h) | 12 |
| Pulse Test (2 h) | 10 |
| Pulse Test (4 h) | 13 |
| Pulse Test (6 h) | 10 |
| Pulse Test (8 h) | 15 |
| Pulse Test (10 h) | 11 |
| Pulse Test (12 h) | 13 |
| Pulse Test (14 h) | 13 |
| Pulse Test (16 h) | 12 |
| Pulse Test (18 h) | 10 |
| Pulse Test (20 h) | 10 |
| 4/18 | 10 |
| 6.5/18 | 13 |
| 9/18 | 10 |
| Adaptive | 8 |

### 5.6 Imaging and analysis

Imaging was conducted using a AxioZoom V16 fluorescence microscope from Zeiss with a Objective Apo Z 1,5x/0,37 at a total magnifiaction of 10.5 over the whole time span of an experiment. Final images were taken using the PlanApo Z 0.5x objective at a total magnification of 6.3. As software for the acquisition we used Zen Blue 3.0. Pictures were than analyzed using a row of Python 3.11 scripts and Ilastik, a machine learning based image segmentation tool, trained on time spaced frames of randomly picked colonies[43]. All time-lapse data shown were segmented with the same trained model, assigning discrete labels for sensitive cells, resistant cells and background. Final endpoint images used for extrapolation were segmented with a separate model to ensure accuracy. For visualisation purposes only, raw experimental images were processed in Fiji to adjust the display lookup table and colormap. Radial kymographs were generated from segmented image stacks. For each colony, the centroid was determined and radial lines were sampled from the centroid to the border. Kymographs were computed over an angular window of ± 2 ° with steps of 0.5 ° and combined by taking the pixel-wise minimum across angles. A 3 x 1 median filter (time x space) was applied for visualisation only.

### 5.7 Clonal Tracking

Clone trajectories were filtered to restrict the analysis to comparably emerging and reliably tracked clones. Only clones whose first detection occurred between frames 50 and 100 were included, excluding early-appearing clones and late detections with insufficient temporal coverage. From this subset, clones were required to enter a defined spatial band relative to the colony front at least once during their lifetime, with distances constrained between 2 and 30 pixels. This criterion restricted the analysis to clones originating within the actively growing region of the colony and excluded both central and edge-contact artifacts. Clones were further required to be tracked for a minimum of three consecutive frames.

Distances to the colony edge were converted to physical units using a scale factor of 8.648 µm per pixel. Clones were classified based on their interaction with the colony front: a breach was defined as reaching a distance of ≤ 20 µm from the colony edge at least once. Following a breach, clones were classified as *escaped* if they did not subsequently retreat beyond 80 µm from the colony edge, whereas clones that never breached or retreated beyond this threshold were classified as *confined*. For visualization purposes only, trajectory traces were smoothed using a Savitzky-Golay filter; all classification decisions were based on unsmoothed data.

### 5.8 Time-to-progression and extrapolation

Time-to-progression was based on a fixed maximal colony area, corresponding to the area a perfectly round colony can occupy in the absence of imaging related truncation. With the field of view used for time resolved imaging, this area was determined to be 71*mm*^2^. Colony- and clone-area trajectories were extrapolated when growth extended beyond the field of view. Under treatment, resistant clones could expand asymmetrically and partially leave the imaging frame, leading to an underestimation of clonal area. For each colony, the first time point at which the colony contour contacted the image border was detected, and the last trusted measurement was defined as the preceding frame. From this frame onward, colony area and total clonal area were extrapolated linearly in physical to a final target area measured from a corresponding endpoint image acquired at a lower magnification. The endpoint was assigned to a time point one frame after the last time-lapse frame. For the no-treatment condition, colony sizes were instead linearly extrapolated on the growth of the resistant population, which showed stable growth dynamics in the absence of therapy. For time-resolved visualisation of population-level trajectories and resistant fraction the median and interquartile range across colonies were computed per time point. To reduce frame-to-frame variability curves were smoothed using a centered rolling median filter (Window size: 4.5 h). Smoothing was applied for display purposes only and did not affect statistical analyses or classification criteria.

### 5.9 Simulation

We built a custom 2D lattice simulation consisting of sensitive and resistant cells interacting via a nutrient layer on a 200 × 200 grid, where the cells have periodic boundary conditions and the nutrients have Dirichlet boundary conditions, leading to a constant influx of nutrients from the boundary. The simulation starts with a single sensitive cell in the center pixel, from which we pre-grow the colony until it reaches a size similar to that observed in the first experimental image.

Space is divided into a 2D grid of demes, each containing a mixture of sensitive cells *S* and resistant cells *R* that proliferate at rate *α*, mutate stochastically (ℳ_*µ*_) and disperse with an effective diffusion coefficient *D*. To emulate the growth-dependent mutation and dispersal of our experiments, these processes are only applied to the actively growing per-deme populations of sensitive and resistant cells *S*_d_ = *αξNS* and *R*_d_ = *αNR*, respectively:

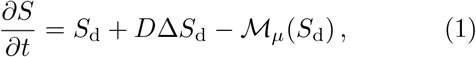

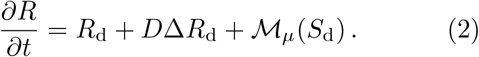

Additionally, dispersal only occurs after a defined population threshold is reached in a deme. This threshold is cell type specific. Experimentally observed delays in treatment effects on growth rate are implemented via the therapy induced growth rate modulation *ξ* ∈ [0, 1], exhibiting an on-delay *δ*_on_, overshoot phase *δ*_over_, lag-delay *δ*_lag_, and off-delay *δ*_off_. Note that the overshoot phase is part of the lag phase (see Supplementary Material). Stochastic mutations are randomly sampled from a Poisson distribution ℳ_*µ*_ defined by the expectation value *µ* · *S*_d_, with *µ* being a mutation prefactor and then divided with a scaling factor to convert the sampled number to the scale used in the simulation.

Importantly, competition is solely mediated via nutrient limitation without introducing a local carrying capacity. Nutrients are modeled as a freely diffusing consumable resource with the diffusion coefficient *D*_N_:

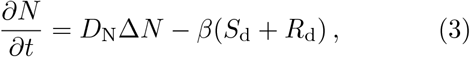

with *β* being the consumption rate of nutrients. If the defined TTP size is not reached, the end point of the simulation is used as TTP value. A description of the fitting of the model parameters to the experiments, is provided in the Supplementary Material.

All median trajectory plots of simulation data are plotted with a rolling average of window size 2.5 h. The area growth rate and front growth rates additionally had a rolling average of the same window size applied during calculation.

### 5.10 Nutrient threshold

The nutrient thresholds *N*_low_ and *N*_high_ which define the position of λ_low_ and λ_high_ are defined as the values of a sigmoidal function at the position where an linear function with the slope of the sigmoid function at 0 intersects 0 and 1 (see Supplementary Fig. 11). With this 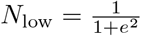 and 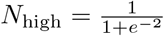.

## 6 Data availability

The source and processed data underlying the figures are available at https://figshare.com/articles/dataset/Appold_Citak_2026/31356370. The analysis code used to generate the figures is available at https://github.com/KayserLab/Appold_Citak_2026.git. Raw imaging datasets are too large for public deposition but are available from the corresponding author upon reasonable request.

## 7 Code availability

Custom code used for the simulations is available at https://github.com/KayserLab/Appold_Citak_2026.git. The repository contains all scripts necessary to reproduce the simulation results and figures presented in this study.

## Acknowledgments

The authors thank S. Aif, M. Eiche, E. Fischer, I. Schneider, and L. Strampe for valuable discussions. We are extremely grateful for the support of J. Guck and his entire division.

## Author contributions

N.A., T.C., and J.K. conceived and designed the study. N.A. and A.P. performed experiments. T.C. performed simulations. N.A., T.C., and A.P. analyzed the data. N.A., T.C., and J.K. wrote the manuscript.

## References

1. Bukowski, K., Kciuk, M. & Kontek, R. Mechanisms of multidrug resistance in cancer chemotherapy. International journal of molecular sciences 21, 3233 (2020).

2. Ingham, J., Ruan, J.-L. & Coelho, M. A. Breaking barriers: we need a multidisciplinary approach to tackle cancer drug resistance. BJC reports 3, 11 (2025).

3. Murray, C. J. et al. Global burden of bacterial antimicrobial resistance in 2019: a systematic analysis. The lancet 399, 629–655 (2022).

4. Naghavi, M. et al. Global burden of bacterial antimicrobial resistance 1990–2021: a systematic analysis with forecasts to 2050. The Lancet 404, 1199–1226 (2024).

5. King, E. S. et al. Fitness seascapes are necessary for realistic modeling of the evolutionary response to drug therapy. Science Advances 11, eadv1268 (2025).

6. Stewart, P. S. & Franklin, M. J. Physiological heterogeneity in biofilms. Nature reviews microbiology 6, 199–210 (2008).

7. Hallatschek, O. & Nelson, D. R. Gene surfing in expanding populations. Theoretical population biology 73, 158–170 (2008).

8. Edmonds, C. A., Lillie, A. S. & Cavalli-Sforza, L. L. Mutations arising in the wave front of an expanding population. Proceedings of the National Academy of Sciences 101, 975–979 (2004).

9. Hallatschek, O. & Nelson, D. R. Life at the front of an expanding population. Evolution 64, 193–206 (2010).

10. Korolev, K. S. et al. Selective sweeps in growing microbial colonies. Physical biology 9, 026008 (2012).

11. Gralka, M. et al. Allele surfing promotes microbial adaptation from standing variation. Ecology letters 19, 889–898 (2016).

12. Kayser, J., Schreck, C. F., Yu, Q., Gralka, M. & Hallatschek, O. Emergence of evolutionary driving forces in pattern-forming microbial populations. Philosophical Transactions of the Royal Society B: Biological Sciences 373, 20170106 (2018).

13. Kayser, J., Schreck, C. F., Gralka, M., Fusco, D. & Hallatschek, O. Collective motion conceals fitness differences in crowded cellular populations. Nature ecology & evolution 3, 125–134 (2019).

14. Giometto, A., Nelson, D. R. & Murray, A. W. Physical interactions reduce the power of natural selection in growing yeast colonies. Proceedings of the National Academy of Sciences 115, 11448–11453 (2018).

15. Fusco, D., Gralka, M., Kayser, J., Anderson, A. & Hallatschek, O. Excess of mutational jackpot events in expanding populations revealed by spatial Luria–Delbrück experiments. Nature communications 7, 12760 (2016).

16. Aif, S., Appold, N., Kampman, L., Hallatschek, O. & Kayser, J. Evolutionary rescue of resistant mutants is governed by a balance between radial expansion and selection in compact populations. Nature communications 13, 7916 (2022).

17. Giometto, A., Nelson, D. R. & Murray, A. W. Antagonism between killer yeast strains as an experimental model for biological nucleation dynamics. Elife 10, e62932 (2021).

18. Lavrentovich, M. O., Wahl, M. E., Nelson, D. R. & Murray, A. W. Spatially constrained growth enhances conversional meltdown. Biophysical journal 110, 2800–2808 (2016).

19. Ascensao, J. A. & Desai, M. M. Experimental evolution in an era of molecular manipulation. Nature Reviews Genetics, 1–15 (2025).

20. Lomakin, A. et al. Spatial genomics maps the structure, nature and evolution of cancer clones. Nature 611, 594–602 (2022).

21. Seferbekova, Z., Lomakin, A., Yates, L. R. & Gerstung, M. Spatial biology of cancer evolution. Nature Reviews Genetics 24, 295–313 (2023).

22. Hallatschek, O. et al. Proliferating active matter. Nature Reviews Physics 5, 407–419 (2023).

23. Schreck, C. F. et al. Impact of crowding on the diversity of expanding populations. Proceedings of the National Academy of Sciences 120, e2208361120 (2023).

24. Zhao, Y. et al. Selection of metastasis competent subclones in the tumour interior. Nature ecology & evolution 5, 1033–1045 (2021).

25. Gatenby, R. A., Brown, J. & Vincent, T. Lessons from applied ecology: cancer control using an evolutionary double bind. Cancer research 69, 7499–7502 (2009).

26. Gatenby, R. A. & Brown, J. S. Integrating evolutionary dynamics into cancer therapy. Nature reviews Clinical oncology 17, 675–686 (2020).

27. Enriquez-Navas, P. M. et al. Exploiting evolutionary principles to prolong tumor control in preclinical models of breast cancer. Science translational medicine 8, 327ra24–327ra24 (2016).

28. Zhang, J., Cunningham, J. J., Brown, J. S. & Gatenby, R. A. Integrating evolutionary dynamics into treatment of metastatic castrate-resistant prostate cancer. Nature communications 8, 1816 (2017).

29. Kim, E., Brown, J. S., Eroglu, Z. & Anderson, A. R. Adaptive therapy for metastatic melanoma: predictions from patient calibrated mathematical models. Cancers 13, 823 (2021).

30. Hockings, H. et al. Adaptive therapy achieves long-term control of chemotherapy resistance in high grade ovarian cancer. bioRxiv (2023).

31. Brady-Nicholls, R. & Enderling, H. Range-bounded adaptive therapy in metastatic prostate cancer. Cancers 14, 5319 (2022).

32. Weaver, D. T., King, E. S., Maltas, J. & Scott, J. G. Reinforcement learning informs optimal treatment strategies to limit antibiotic resistance. Proceedings of the National Academy of Sciences 121, e2303165121 (2024).

33. Das Thakur, M. et al. Modelling vemurafenib resistance in melanoma reveals a strategy to forestall drug resistance. Nature 494, 251–255 (2013).

34. Algazi, A. P. et al. Continuous versus intermittent BRAF and MEK inhibition in patients with BRAF-mutated melanoma: a randomized phase 2 trial. Nature medicine 26, 1564–1568 (2020).

35. Maltas, J. et al. Drug dependence in cancer is exploitable by optimally constructed treatment holidays. Nature ecology & evolution 8, 147–162 (2024).

36. Bacevic, K. et al. Spatial competition constrains resistance to targeted cancer therapy. Nature communications 8, 1995 (2017).

37. Noble, R. et al. Spatial structure governs the mode of tumour evolution. Nature ecology & evolution 6, 207–217 (2022).

38. Lewinsohn, M. A., Bedford, T., Müller, N. F. & Feder, A. F. State-dependent evolutionary models reveal modes of solid tumour growth. Nature Ecology & Evolution 7, 581–596 (2023).

39. Strobl, M. A. et al. Spatial structure impacts adaptive therapy by shaping intra-tumoral competition. Communications medicine 2, 46 (2022).

40. Gallaher, J. A., Enriquez-Navas, P. M., Luddy, K. A., Gatenby, R. A. & Anderson, A. R. Spatial heterogeneity and evolutionary dynamics modulate time to recurrence in continuous and adaptive cancer therapies. Cancer research 78, 2127–2139 (2018).

41. Gallaher, J. et al. Intermetastatic and intrametastatic heterogeneity shapes adaptive therapy cycling dynamics. Cancer Research 83, 2775–2789 (2023).

42. Wahl, M. E. & Murray, A. W. Multicellularity makes somatic differentiation evolutionarily stable. Proceedings of the National Academy of Sciences 113, 8362–8367 (2016).

43. Berg, S. et al. Ilastik: interactive machine learning for (bio) image analysis. Nature methods 16, 1226–1232 (2019).

