## Supplementary Information for "Spatial resource dynamics control resistance escape"

### Simulation: Fitting

The treatment efficacy $E$ increases linearly with time from zero to one during treatment, reaching full effect after 250 simulation steps (12.5 h). If the treatment ends before full effect is reached $E$ continuous to increase for another 30 simulation steps (1.5 h). After treatment ends $E$ has a 220 simulations steps (11 h) lag phase including the overshoot time before it decreases linearly back to zero in 240 simulation steps (12 h). In case that treatment starts again before zero is reached an immediate switch occurs to the treatment on dynamics. These delays are determined directly from the experiment by analyzing the radial growth over time (Supplementary Fig. 3a) for a 7/18 intermittent and 14 h pulse experiment. Example effective growth rates ($\xi = 1 - E$) are shown in Supplementary Fig. 7. The growth rate $\alpha$ is calculated from the doubling time $\tau_2$ of the sensitive cell population determined via a plate reader experiment using:

$$\alpha = \frac{\log(2)}{\tau_2}. \tag{1}$$

The cell diffusion coefficient $D$, the nutrient diffusion coefficient $D_N$ and the consumption rate $\beta$ are fitted together using the Nelder-Mead method. The value we minimize is the sum of two normalized mean squared errors (NMSE), each focused an a different system dynamic - but not independent. The first part is calculated by comparing the radial expansion of the experiment for 13 experiment replicates in the case of no treatment to the simulation (Supplementary Fig. 3b). We calculate the NMSE for each replicate and then average over the replicates giving us the first part of the sum. The second part is calculated from the time it takes for a clone to start growing again after treatment starts over the distance to the front (experiment data from Fig. 2). We choose four distances (3 px, 7 px, 11 px, 15 px) and initialize the simulation with exactly one mutated cell at the corresponding distance along the cardinal directions, simulate treatment and compare the time it takes for the mutated population to reach the next pixel closer to the front to the time measured from the experiment (Supplementary Fig. 3c). For this we again calculate the NMSE. Taken the result of this fit we next fit the mutation prefactor by minimizing the squared difference between the number of mutation events in the simulation and the experiment in the case of continuous treatment using the Nelder-Mead method. Finally the mutation scaling is fitted to the breakout probability of a clone. For this we analyzed the number of colonies which had at least one breakout over a variety of pulse durations. For each pulse duration we simulated twice the number of colonies as we have in the experiment (Tab. 1), calculated the breakout probability at the end point and calculated the squared distance to the experimental value, which is then used as the minimization value. The resulting breakout probability for experiment vs simulation (75 simulation replicates per pulse duration) is shown in Supplementary Fig. 3d. The final parameter set is available in the code repository.

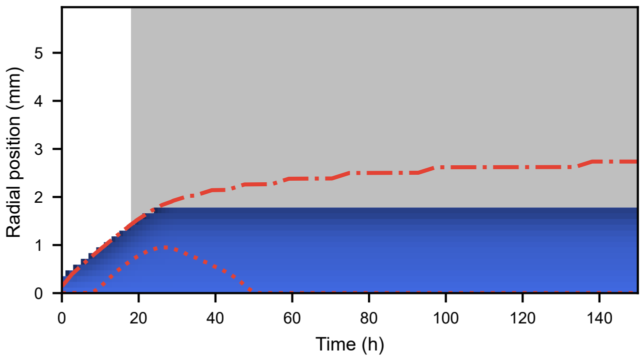


**Supplementary Figure** 1: Example of how the nutrient gradient evolves under treatment if no mutations occur.

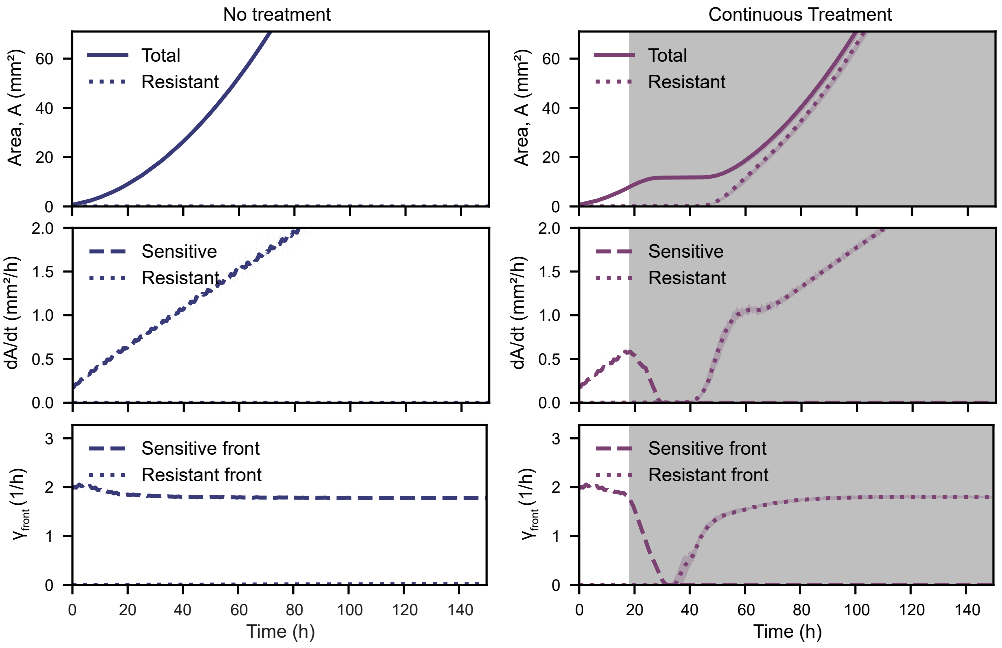


**Supplementary Figure** 2: Total and resistant area, sensitive and resistant area growth rates and front cell growth speeds and for the no treatment and continuous treatment scenarios. The lines are the median over n = 20 colonies, with the shaded area being the IQR. Gray shading indicates treatment.

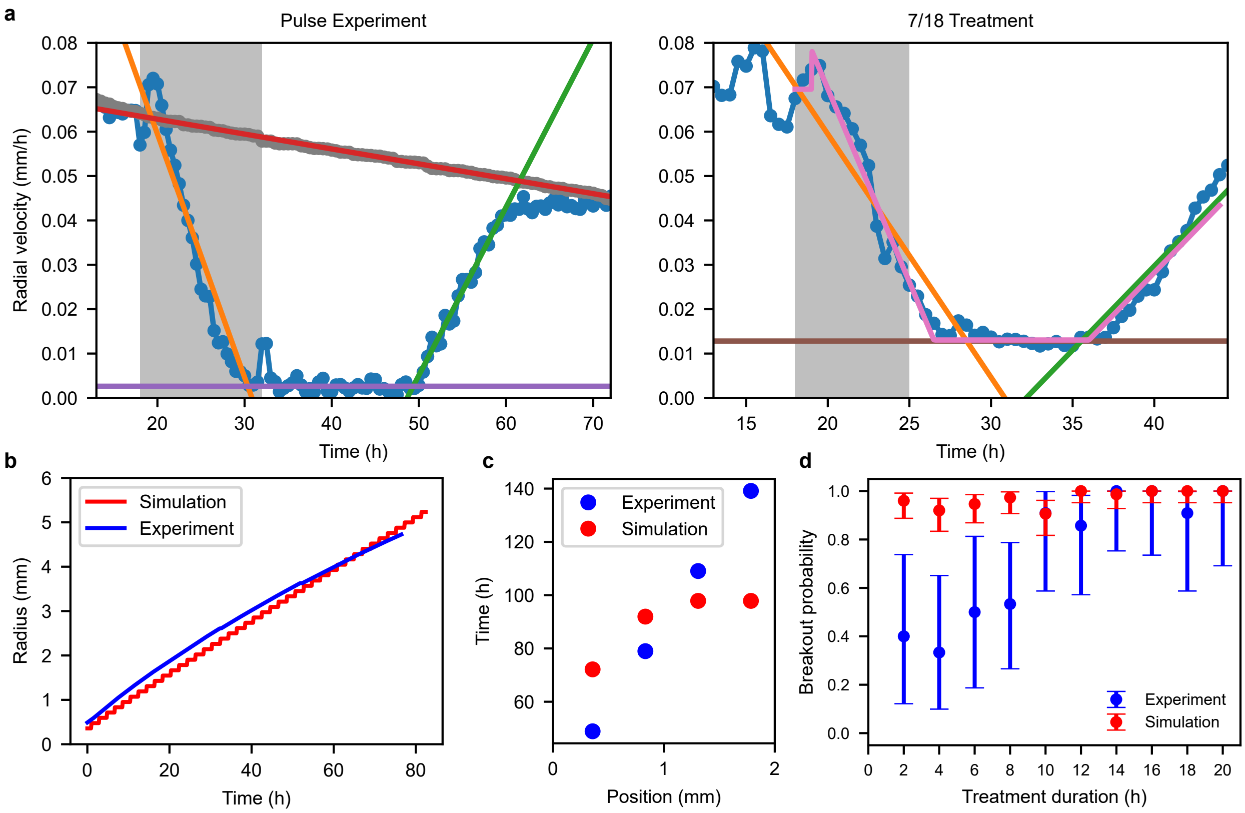


**Supplementary Figure** 3: **a** Treatment delays fit: The orange line in the left panel from treatment start until it intersects with the purple line gives the duration of the $\delta_{\mathrm{on}}$ delay. The x axis difference between the intersection of the green line with the purple line and the red line gives the $\delta_{\mathrm{off}}$ delay, the red line being the maximum radial velocity at that time point taken from a no treatment experiment. The right panel gives the $\delta_{\mathrm{over}}$ and $\delta_{\mathrm{lag}}$ delays. The lag delay is the x axis difference from the treatment end point until pink line starts to increase again, while the overshoot delay is the short duration after treatment ends it still takes for the pink line to reach its minimum. Gray shading indicates treatment. **b** Radial expansion fit in the no treatment scenario. **c** Time it takes a resistant clone to start growing again after treatment starts at 18 h dependent on distance from colony front. Treatment is continuous dose. **d** Probability for a colony to have at least one breakout for the simulation and the experiment, dependent the duration of a single treatment pulse. The error bars are the Clopper-Pearson Intervall. The sample size of the simulation is n = 75, while the experimental sample size is the same as in Fig. 2.

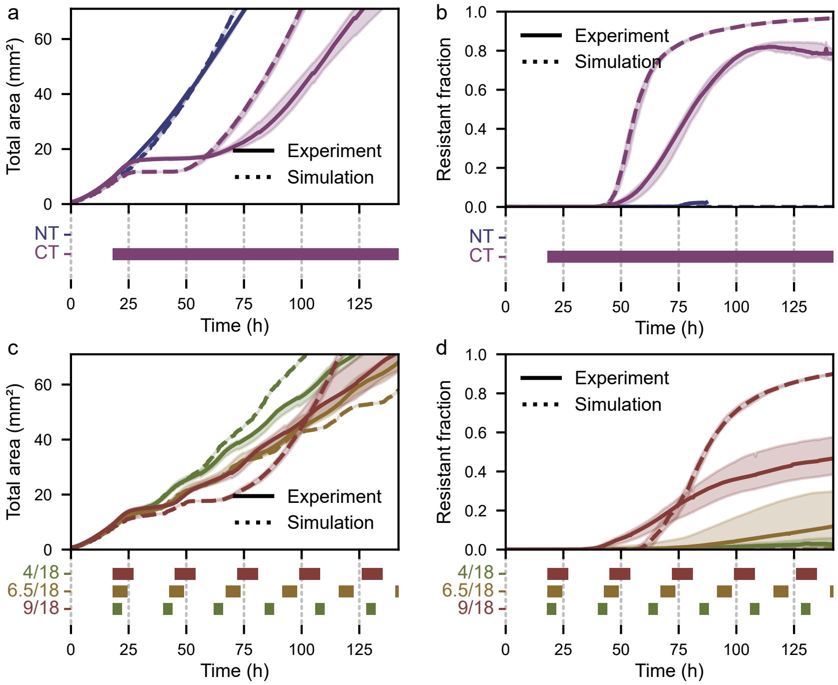


**Supplementary Figure** 4: **a - d** Direct comparison between the simulation and the experiment in total area and resistant fraction for the treatment schedules shown in Fig 5. Lines are the median, shaded areas are the IQR. The sample size for the simulation is $n = 20$, the experiment sample size is the same as in Fig. 5. The colored bars indicate treatment.

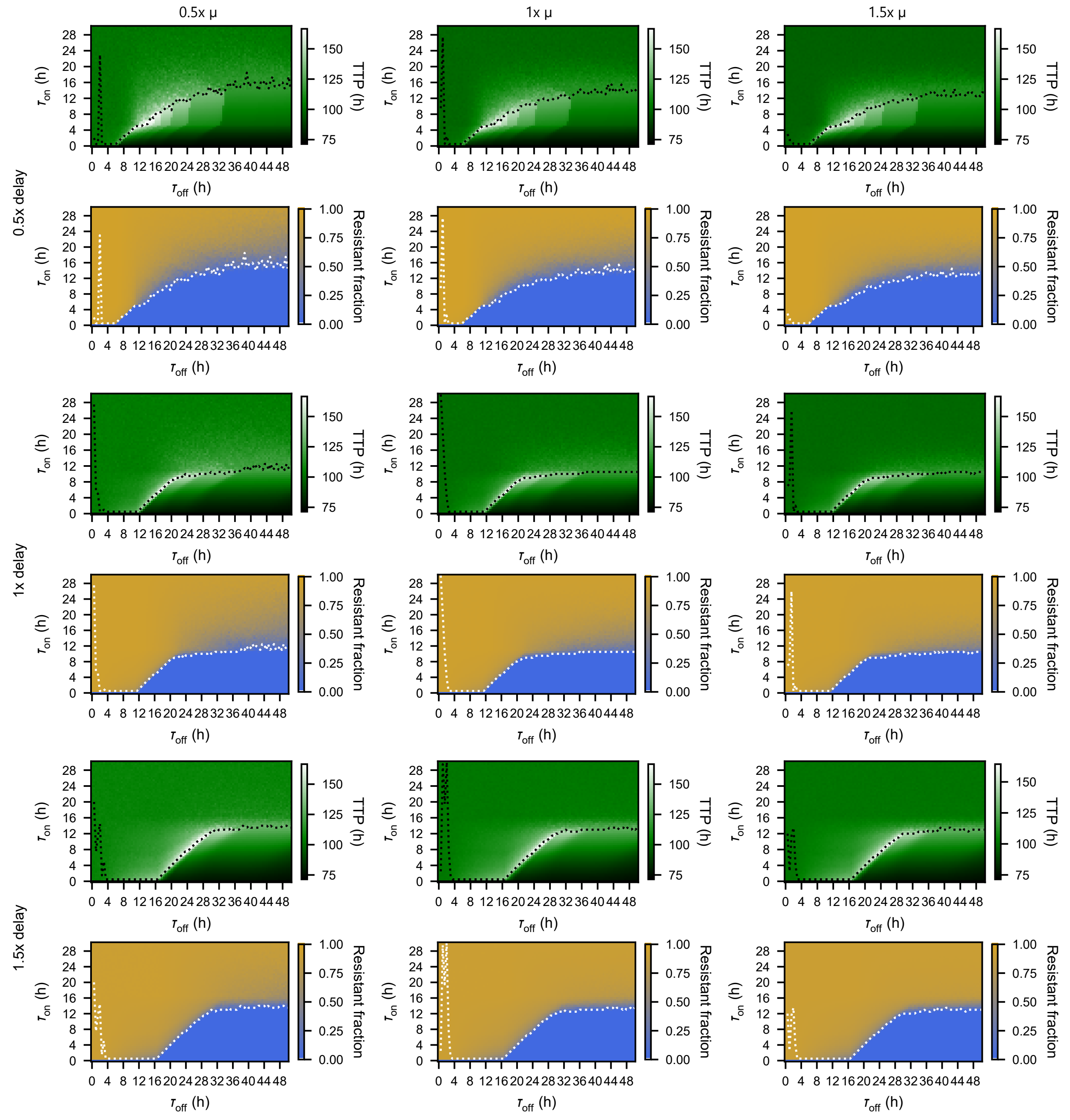


**Supplementary Figure** 5: Full parameter sweep plots for the sweep shown in Fig. 4a,b as well as sweeps with 0.5x and 1.5x the mutation prefactor and total delay duration. Each pixel is the median of 20 replicates.

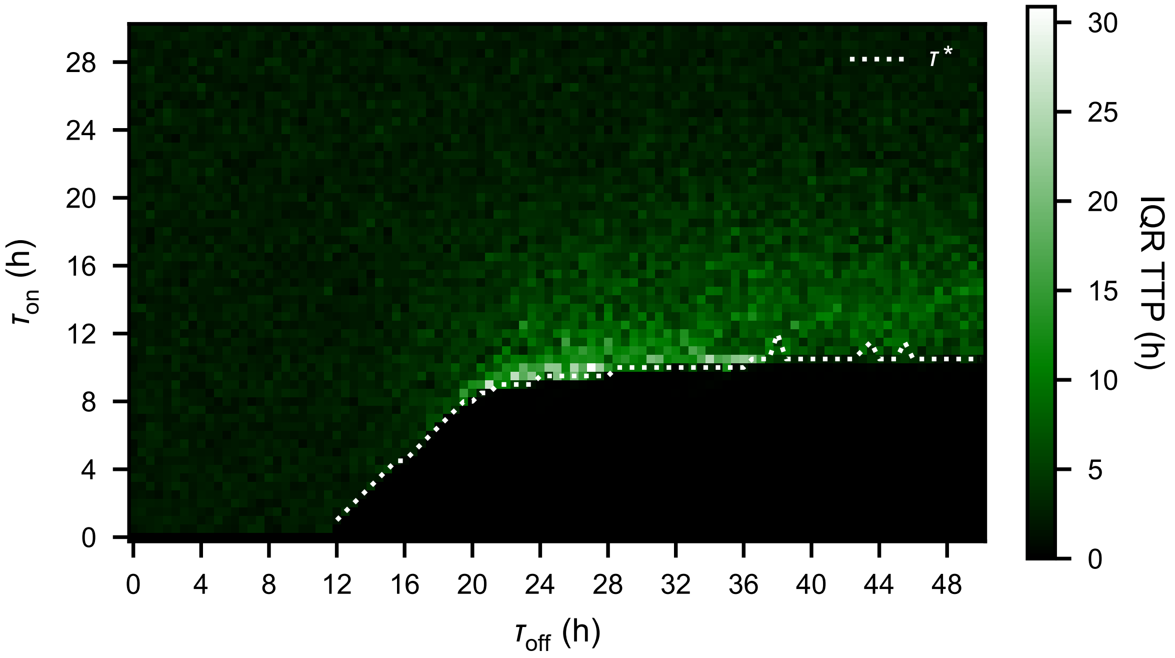


**Supplementary Figure** 6: IQR for the full parameter sweep of which parts are shown in Fig. 4a,b.

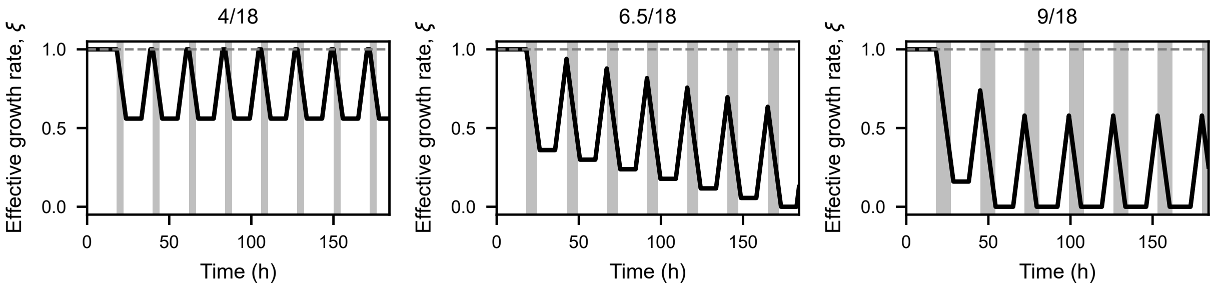


**Supplementary Figure** 7: Simulated effective growth rates $\xi$ due to treatment for the intermittent treatments shown in Fig. 4. Gray shading indicates treatment.

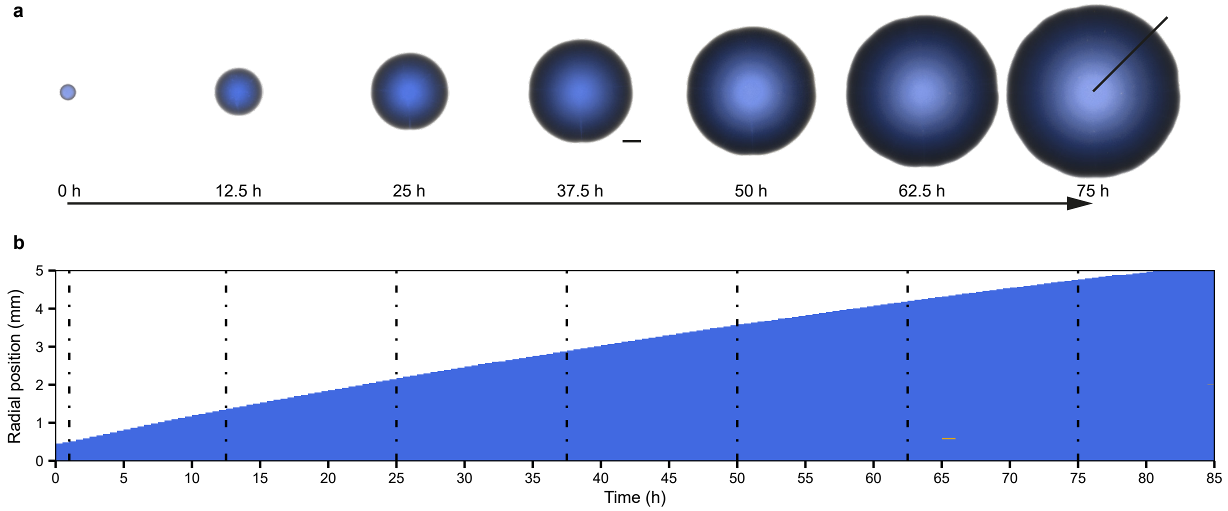


**Supplementary Figure** 8: **a** Representative time-lapse images of an no treatment experiment. Imaging starts ($t$ = 0 h) 24 h after seeding. Time points are at 1, 12.5, 25, 37.5, 50 and 62.5 h after imaging start. Scale bar indicates 1 mm. **b** Radial kymographs extracted from the colony depicted in **a** (black radial at 45 °). Dash-dotted lines correspond to the time points shown in (**a**)

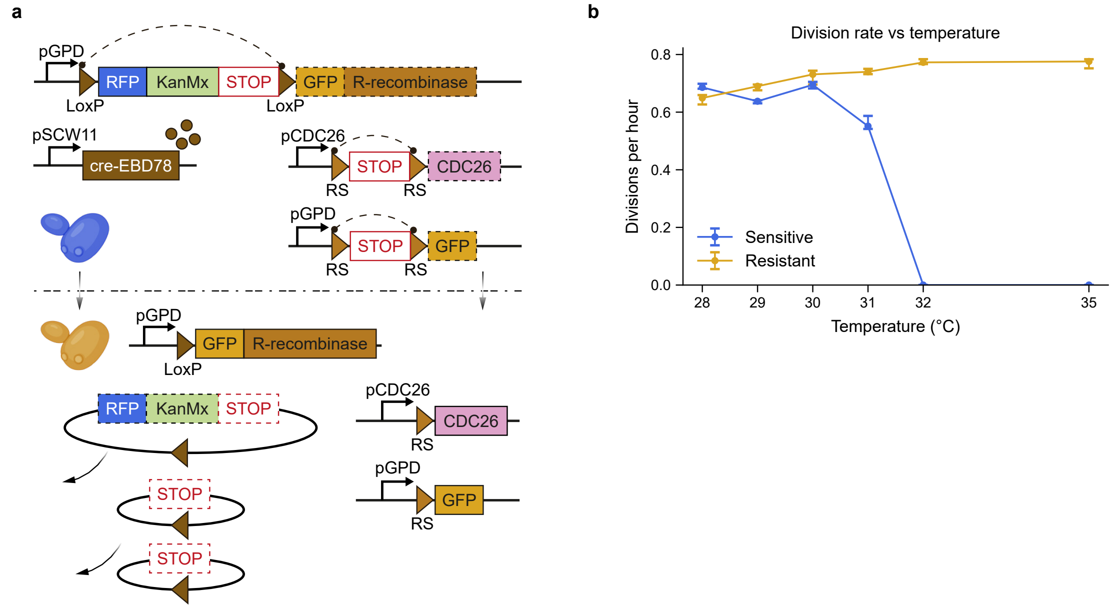


**Supplementary Figure** 9: **a** Detailed schematic of the yNA16 genetic construct showing the synthetic mutation enabling the switch from thermo-sensitive to thermo-resistant cells via activation of an R-recombinase cascade through Cre-Lox recombination . **b** Median divisions rates of sensitive and resistant cells in liquid culture at the indicated temperatures. Whiskers show interquartile range.

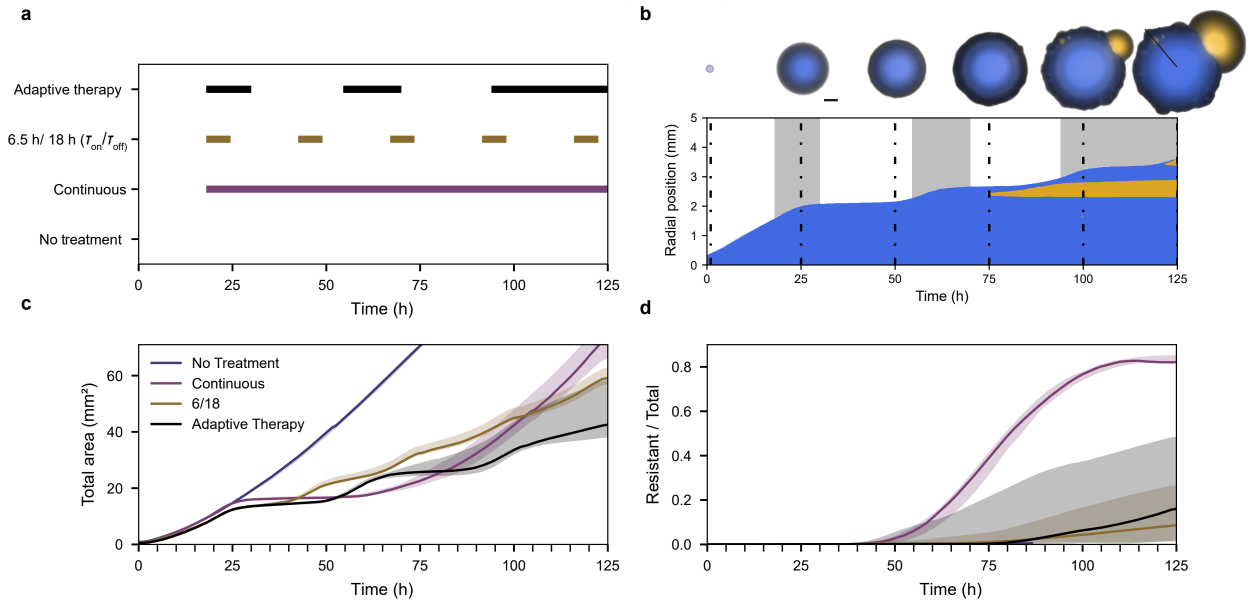


**Supplementary Figure** 10: **a** Treatment schedules used for adaptive therapy, 6.5 h/ 18 h ($\tau_{\mathrm{on}}/\tau_{\mathrm{off}}$), continuous therapy and no treatment. **b** Time lapse images and radial kymograph (320 °) of a representative colony under adaptive therapy. **c** Total population size over time under the indicated treatment schedules. Lines indicate the median; shaded areas denote the interquartile range **d** MFraction od resistant cells over time under the indicated treatment schedules. Lines indicate the median; shaded areas denote the interquartile range

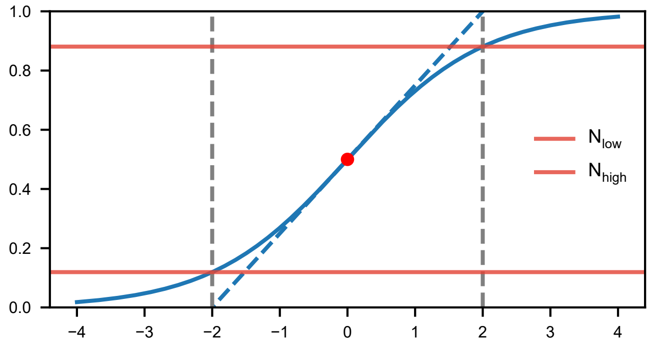


**Supplementary Figure** 11: Definition of the nutrient threshold $N_{\mathrm{low}}$ and $N_{\mathrm{high}}$, which define the position of $\lambda_{\mathrm{low}}$ and $\lambda_{\mathrm{high}}$. The blue line is a sigmoidal function and the dashed blue line is a linear function with the slope of the sigmoid function at the red dot. The grey lines indicate the intersection of the linear function with 0 and 1.

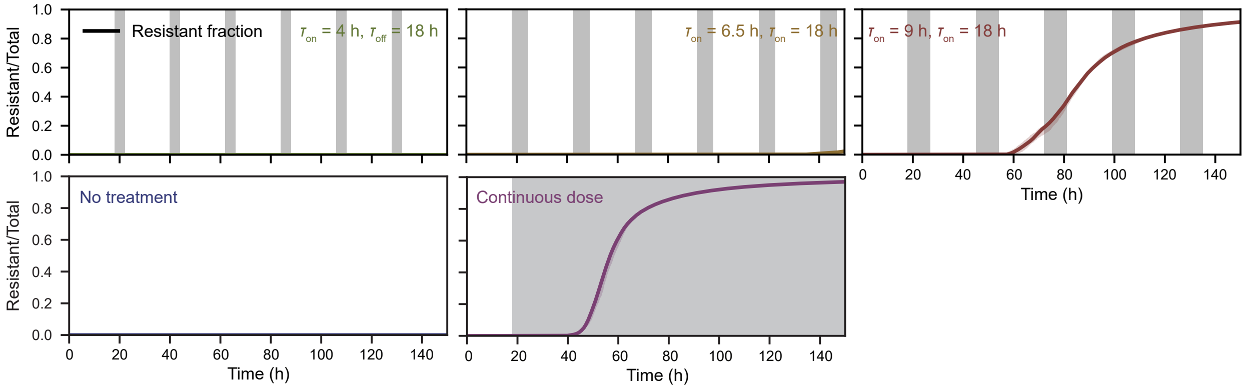


**Supplementary Figure** 12: Median resistant fraction (n = 20) with their respective IQR for the treatments shown in Fig. 5. Gray shading indicates treatment.
